# Variability in an effector gene promoter of a necrotrophic fungal pathogen dictates epistasis and effector-triggered susceptibility in wheat

**DOI:** 10.1101/2021.07.28.454099

**Authors:** Evan John, Silke Jacques, Huyen T. T. Phan, Lifang Liu, Danilo Pereira, Daniel Croll, Karam B. Singh, Richard P. Oliver, Kar-Chun Tan

## Abstract

The fungus *Parastagonospora nodorum* uses proteinaceous necrotrophic effectors (NEs) to induce tissue necrosis on wheat leaves during infection, leading to the symptoms of septoria nodorum blotch (SNB). The NEs Tox1 and Tox3 induce necrosis on wheat possessing the dominant susceptibility genes *Snn1* and *Snn3B1/Snn3D1*, respectively. We previously observed that *Tox1* is epistatic to the expression of *Tox3* and a quantitative trait locus (QTL) on chromosome 2A that contributes to SNB resistance/susceptibility. The expression of *Tox1* is significantly higher in the Australian strain SN15 compared to the American strain SN4. Inspection of the *Tox1* promoter region revealed a 401 bp promoter genetic element in SN4 positioned 267 bp upstream of the start codon that is absent in SN15, called PE401. Analysis of the world-wide *P. nodorum* population revealed that a high proportion of Northern Hemisphere isolates possess PE401 whereas the opposite was observed in the Southern Hemisphere. The presence of PE401 ablates the epistatic effect of *Tox1* on the contribution of the SNB 2A QTL but not *Tox3*. PE401 was introduced into the *Tox1* promoter regulatory region in SN15 to test for direct regulatory roles. *Tox1* expression was markedly reduced in the presence of PE401. This suggests a repressor molecule(s) binds PE401 and inhibits *Tox1* transcription. Infection assays also demonstrated that *P. nodorum* which lacks PE401 is more pathogenic on *Snn1* wheat varieties than *P. nodorum* carrying PE401. An infection competition assay between *P. nodorum* isogenic strains with and without PE401 indicated that the higher *Tox1*-expressing strain rescued the reduced virulence of the lower *Tox1*-expressing strain on *Snn1* wheat. Our study demonstrated that *Tox1* exhibits both ‘selfish’ and ‘altruistic’ characteristics. This offers an insight into a ’NE arms race’ that is occurring within the *P. nodorum* population. The importance of PE401 in breeding for SNB resistance in wheat is discussed.

**Author summary:** Breeding for durable resistance to fungal diseases in crops is a continual challenge for crop breeders. Fungal pathogens evolve ways to overcome host resistance by masking themselves through effector evolution and evasion of broad-spectrum defense responses. Association studies on mapping populations infected by isolate mixtures are often used by researchers to seek out novel sources of genetic resistance. Disease resistance quantitative trait loci (QTL) are often minor or inconsistent across environments. This is a particular problem with septoria diseases of cereals such as septoria nodorum blotch (SNB) of wheat caused by *Parastagonospora nodorum*. The fungus uses a suite of necrotrophic effectors (NEs) to cause SNB. We characterised a genetic element, called PE401, in the promoter of the major NE gene *Tox1*, which is present in some *P. nodorum* isolates. PE401 functions as a transcriptional repressor of *Tox1* and exerts epistatic control on another major SNB resistance QTL in the host. In the context of crop protection, constant surveillance of the pathogen population for the frequency of PE401 in conjunction with NE diversity will enable agronomists to provide the best advice to growers on which wheat varieties can be tailored to provide optimal SNB resistance to regional pathogen population genotypes.

## Introduction

Breeding for effective resistance to fungal diseases in crops is a continual challenge. Fungal pathogens, namely biotrophs and hemibiotrophs, evolve ways to overcome host resistance by masking themselves to evade pathogen-triggered immunity. This can occur through mutations in *Avr-*effector genes, leading to the loss of recognition and the subsequent breakdown of host resistance that ultimately results in ‘boom and bust’ cycles (1). This is manifested through the mass adoption of a resistant cultivar (boom). The pathogen population exposed to the resistant gene will then adapt by selecting favourable mutations in the *Avr* gene that leads to a loss of host recognition and resistance (bust). Notable examples occur in the rust and blackleg pathosystems (2, 3). In contrast, necrotrophic fungal pathogens belonging to the Pleosporales such as the tan spot of wheat fungus *Pyrenophora tritici-repentis* (*Ptr*) and the septoria nodorum blotch of wheat fungus *Parastanonospora nodorum* secrete host-specific necrotrophic effectors (NEs) to cause tissue necrosis on host plants possessing a matching dominant susceptibility gene. This results in effector-triggered susceptibility, enabling the pathogen to proliferate (4, 5). Improvements in host resistance to necrotrophic fungal pathogens rely on the removal of the host dominant-susceptibility genes (6). However, necrotrophic fungal pathogens often possess multiple NEs that exploit different dominant- susceptibility genes carried by the host (7). Hence, these functional redundancies often require breeders to remove multiple host genes or stack the required number of desirable disease resistance alleles to be effective (8, 9). These situations make it difficult for breeders to provide long-lasting and durable resistance.

*P. nodorum* utilises proteinaceous necrotrophic effectors (NE)s to cause necrosis on host plants with a susceptible genotype, leading to necrosis and SNB of wheat (10–12). Genetic evidence has revealed at least 10 NE-dominant susceptibility gene interactions in the *P*. *nodorum-*wheat pathosystem. These are ToxA-*Tsn1* (13, 14), Tox1-*Snn1* (15, 16), Tox267-*Snn2*/*Snn6*/*Snn7* (17), Tox3-*Snn3B1*/*Snn3D1* (18, 19), Tox4-*Snn4* (20), Tox5-*Snn5* (21) and Tox2A-*Qsnb.cur–2AS1* (22). Of the NEs, genes for ToxA (13), Tox1 (16), Tox3 (18), Tox267 . (17) and Tox5 (21) have been cloned and characterised for their roles in virulence. These NEs are secreted, generally have a high cysteine content and are relatively small in molecular mass (<30kDa). Of the host dominant-susceptibility receptors, the genes for *Tsn1* (14), *Snn1* (15) and *Snn3D1* (19) have been cloned. These genes encode proteins similar to canonical nucleotide-binding leucine-rich repeat resistance gene receptors that typically possess a protein kinase domain coupled with nucleotide-binding leucine-rich repeats or major sperm protein domains. Perturbation of NE-host dominant susceptibility gene interactions, either through the loss of NE or the host receptor, results in ablation of ETS and a drastic reduction of *P. nodorum* virulence on wheat (eg. 13, 16, 18).

SNB resistance is quantitatively inherited (23) and improvements in resistance have been, in part, mediated through the removal of dominant susceptibility genes in the host (6). However, the situation is complex. Multiple NEs expressed by *P. nodorum* afford the pathogen functional redundancies on wheat varieties that possess multiple dominant susceptibility genes (8, 24). For instance, Tan *et al.* (8) demonstrated that the deletion of *ToxA*, *Tox1* and *Tox3* in . *P. nodorum* SN15 did not significantly alter virulence on a diverse collection of modern wheat varieties (8, 9). A further complication in breeding for SNB resistance is that some NE-host gene interactions were not consistently detected in field trials using mapping populations with known susceptibility genes (25, 26). It is intuitive to hypothesise that NE epistasis plays a significant role in shaping the variations observed in host resistance to SNB (Table 1) (27). NE epistasis can be defined as interactions between NE genes where the effect conferred by one is masked by the presence of another (27). NE epistasis in the *P. nodorum-*wheat pathosystem can be divided into two broad regulatory categories. Firstly, the contribution of a NE-host dominant susceptibility gene interaction is suppressed by another competing NE-host gene interaction possibly through host-gene action or cross-talk among NE recognition pathways (eg. 28, 29). Secondly, NE gene repression mediated by the expression of another NE (eg. 22, 30). For the latter, a clear-cut example involves the suppression of the Tox3-*Snn3* interaction in SNB by Tox1-*Snn1* described by Phan *et al.* (22). This was explored as part of a study that mapped SNB on a double-haploid (DH) wheat population derived from two major Australian commercial wheat cultivars. SNB quantitative trait loci (QTL) detected during infection by the Australian *P*. *nodorum* reference isolate SN15 were compared with a *Tox1* deletion mutant (*tox1-6*) (22). This revealed the Tox1-*Snn1* NE-host receptor interaction was epistatic to Tox3- *Snn3B1* and a major SNB QTL located on chromosome 2A in the DH population that confers sensitivity to a novel NE which we coined as Tox2A. Quantitative RT-PCR revealed that *Tox3* expression was significantly higher in *tox1-6* than in the wildtype, which presented a basis for NE epistasis mediated by gene repression (22).

**Table 1.** Epistasis of *P. nodorum* NE-wheat dominant susceptibility gene interactions. *RI, recombinant inbred; DH, doubled haploid; ITMI, International Triticeae Mapping Initiative.

| Interaction suppressed | Suppressing interaction | Isolate | Wheat population* | Ref |
| --- | --- | --- | --- | --- |
| Tox1-*Snn1* | ToxA-*Tsn1* | SN2000, SN4, SN5, SN15 | Sumai 3 x CS-DIC-5B (RI) | (31) |
|  | Tox3-*Snn3B1* | SN4, SN5, SN15 | Sumai 3 x CS-DIC-5B (RI) | (31) |
|  | Tox267-*Snn2/6* | SN4 | ITMI (RI) | (17) |
| Tox267-*Snn2* | ToxA-*Tsn1* and/or Tox3-*Snn3B1* | SN15 | Calingiri x Wyalkatchem (DH) | (22) |
| Tox3-*Snn3* | ToxA-*Tsn1* | SN15, SN1501 | BR34 x Grandin (RI) | (28) |
|  | Tox1-*Snn1* | SN15 | Calingiri x Wyalkatchem (DH) | (22) |
|  | Tox2-*Snn2* | SN15, SN1501 | BR34 x Grandin (RI) | (28) |
|  | Tox5-*Snn5* | SN1501 | Lebsock x PI94749 (DH) | (29) |
|  | Tox267-*Snn2/6* | SN4 | ITMI (RI) | (17) |
| Tox2A-*Qsnb.cur-2AS1* | Tox1-*Snn1* | SN15 | Calingiri x Wyakatchem (DH) | (22; this study) |

The expression of *Tox1* also varies among isolates. As Tox1 production was much higher in SN2000 than both SN4 and SN6, the former was used to enable its detection and functional characterisation (16, 32, 33). This contrasts with other USA isolates such as SN4 and SN6 where *Tox1* expression is low relative to SN2000 (34). A recent study also demonstrated that the Australian *P*. *nodorum* SN15 isolate expressed *Tox1* significantly higher than SN4, which correlated with a greater SNB disease contribution mediated by the Tox1- *Snn1* interaction (31). The mechanisms behind this differential expression remain unexplored however, it was noted that the *Tox1* promoter region was found to be polymorphic between these isolates (31). These observations prompted us to ask several key questions. Firstly, does the genetic polymorphism in the promoter region drive differential *Tox1* expression and affect epistatic outcome of Tox3-*Snn3B1* and Tox2A-*Qsnb.cur–2AS1* interactions. Secondly, what is the possible mechanism of *Tox1* regulation? Thirdly, is there a fitness penalty associated with lower *Tox1* expression in *P. nodorum* isolates? And lastly, what is the global distribution of *Tox1*-expressing *P. nodorum* variants? The study presented here was undertaken to characterise the gene regulatory elements controlling *Tox1* expression in *P*. *nodorum* and explore their role in NE epistasis with end-goals of 1. improving SNB management strategies in wheat growing regions where the disease is prominent and 2. to provide an insight into a mechanism of NE epistasis that may dictate the outcome of fungal-host plant pathosystems.

## Results

### Genetic variability in the *Tox1* promoter region

To determine why both *Tox1-* mediated epistasis and *Tox1* expression differed between. *P. nodorum* isolates, we aligned *Tox1* promoters sequences from a geographically diverse set of 24 *Parastagonospora spp*. genomes (35–37) (Fig 1A). The alignment revealed a 401 bp genetic element that was absent in four isolates within the collection. This element, henceforth called PE401 was absent in the Australian *P*. *nodorum* SN15 and SN2000 isolates that were associated with a higher level of *Tox1* expression (Fig 1A) (16, 31, 34). The size and proximity to the gene within the *Tox1* promoter suggested PE401 was the most plausible variant involved in gene repression, despite the presence of other smaller genetic polymorphisms.

**Fig 1.**
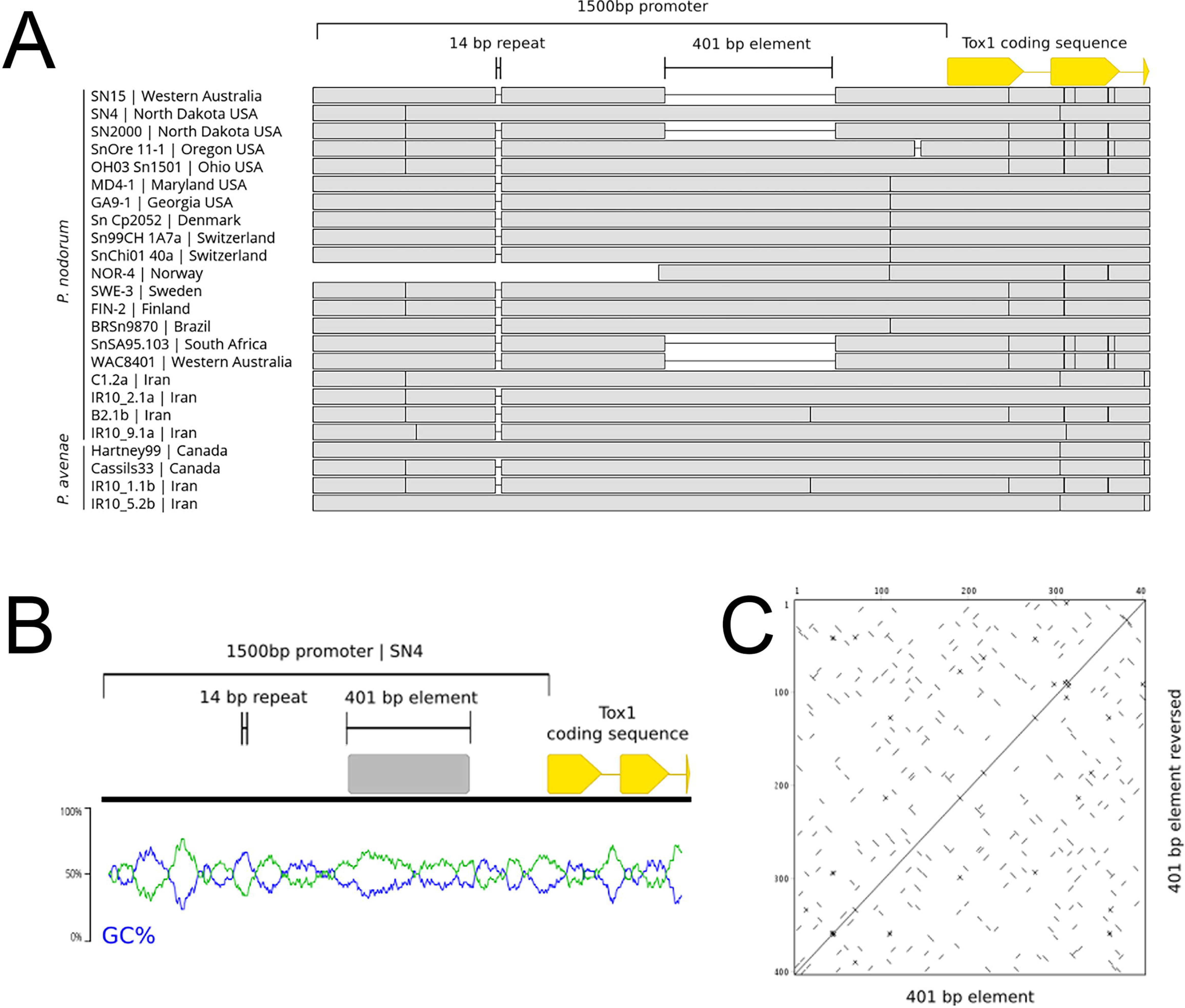
Genetic variation in the *Tox1* promoter region. An alignment highlighting promoter variants in a geographically diverse selection of 24 *Tox1*-positive *Parastagonospora spp.* isolates. Within the 1,500 bp promoter region, the 401 bp PE401 and a 14 bp GTTTTCGGCCGTAT tandem repeat polymorphism are indicated. B. A GC% plot using a 50 bp average sliding window covering the *Tox1* promoter from the USA reference isolate SN4. PE401 has a low GC% relative to directly adjacent promoter regions. C. A dot plot of PE401 using a 5 bp sliding window demonstrating the paucity of tandem/inverted repeat stretches. Genome coverage for of the promoter region of *Tox1* in NOR-4 is incomplete.

PE401 was then examined for features of a mobile genetic element (MGE), as they are known to modulate the expression of proximal fungal genes (38–41). The SN4 promoter sequence was queried against the Repbase and Rfam databases. No matches to characterised MGEs or expressed non-coding elements were observed. BLAST analysis of PE401 identified a single region with 63% identity in the SN15 genome, partially overlapping the annotated gene SNOG_30065 (35). However, the SNOG_30065 predicted peptide sequence did not contain annotated protein domains, nor was the gene expressed under *in vitro* conditions and during host infection (42). Therefore, SNOG_30065 is likely a pseudogene. No further similar matches to PE401 were detected when queried against the NCBI nucleotide and whole-genome shotgun collections, suggesting the sequence was not a repetitive element. Terminal repeats characteristic of fungal MGEs were not identified, although the GC% was low (41.9%) relative to the rest of the SN4 promoter region (48.9%) (43) (Fig 1B and 1C). We concluded that PE401 is unlikely to be a transposable element or functional gene in *P. nodorum*.

### Evidence of regional distribution for PE401 in the global *P. nodorum* population

The distribution of PE401 was determined in a global panel of 489 *Tox1*-containing *P*. *nodorum* isolates. It was observed that isolates carrying PE401 were ubiquitous in Europe (Switzerland, Denmark, Finland, Netherlands and Norway), most sampling sites in North America (USA and Canada) and also in Iran. The latter region was previously proposed as a centre of origin for *P. nodorum* NEs including Tox1 (44, 45). Isolates that lacked PE401 were dominant in the Australian (87% of 179 isolates) and South African (100% of 22 isolates) populations, while 27% of all isolates from North Dakota (USA) and a single isolate from Texas (USA) also lacked PE401 (Fig 2). Therefore, a major shift in the frequency of PE401 was observed between wheat-growing regions worldwide.

**Fig 2.**
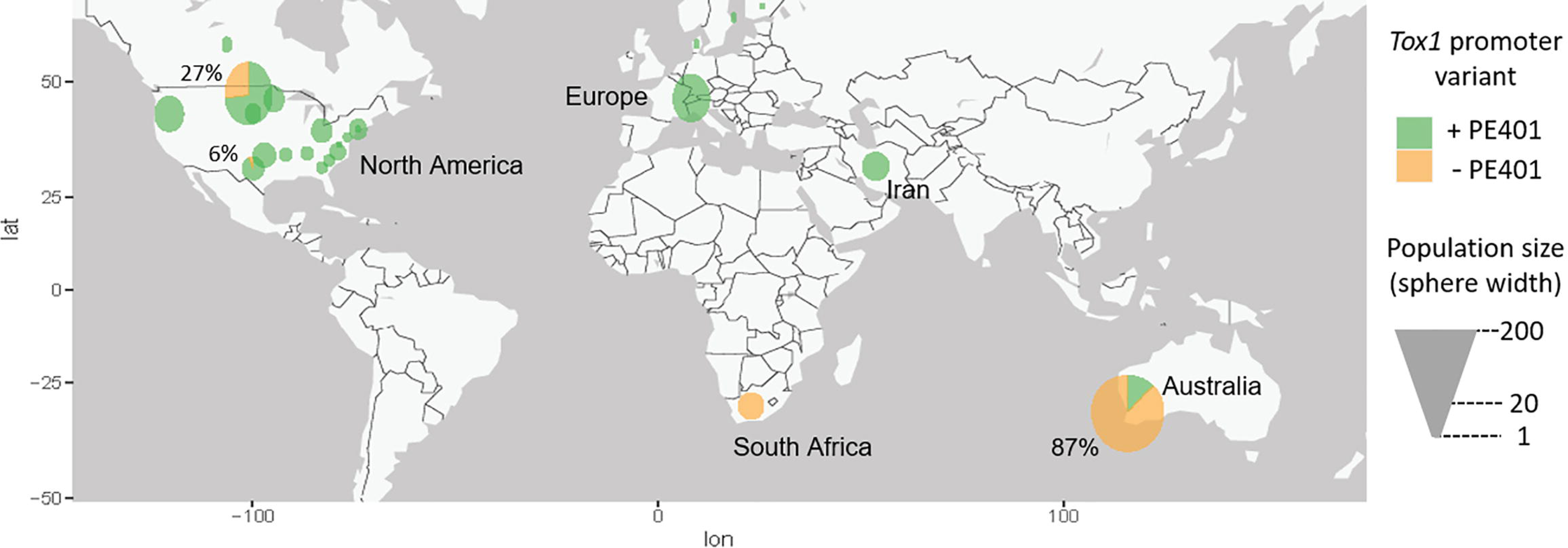
The global distribution of PE401 in the *P. nodorum Tox1* promoter. Presence/absence of PE401 in a global collection of 489 *P. nodorum* isolates. Spheres represent individual populations based on the country of origin or, for the USA and Australia. % values represent the proportion of (-) PE401 isolates where it was identified in that population. Isolate details are provided in S1 File. Image from Google (CA, USA).

### PE401 functions as a repressor of *Tox1* expression

We hypothesised that PE401 dictates *Tox1* expression based on the differential expression profile between SN4 (+PE401) and SN15 (-PE401) carrying the two versions of the promoter (31). Therefore, *Tox1* expression was assessed in SN4, SN15 and three representative Australian *P. nodorum* wildtype isolates with or without PE401 (Fig 3A). Quantitative RT-PCR analysis revealed the two Australian isolates carrying PE401, WAC13443 and WAC13072, shared a low *Tox1* expression profile comparable to SN4. On the other hand, *Tox1* expression was significantly higher in SN15 and WAC13690 which lacked PE401 (Fig 3B). Sequence alignment of the promoter region revealed WAC13443 and WAC13072 does not carry the 14 bp tandem repeat found in SN4, indicating that PE401 was the variant responsible (Fig 3A).

**Fig 3.**
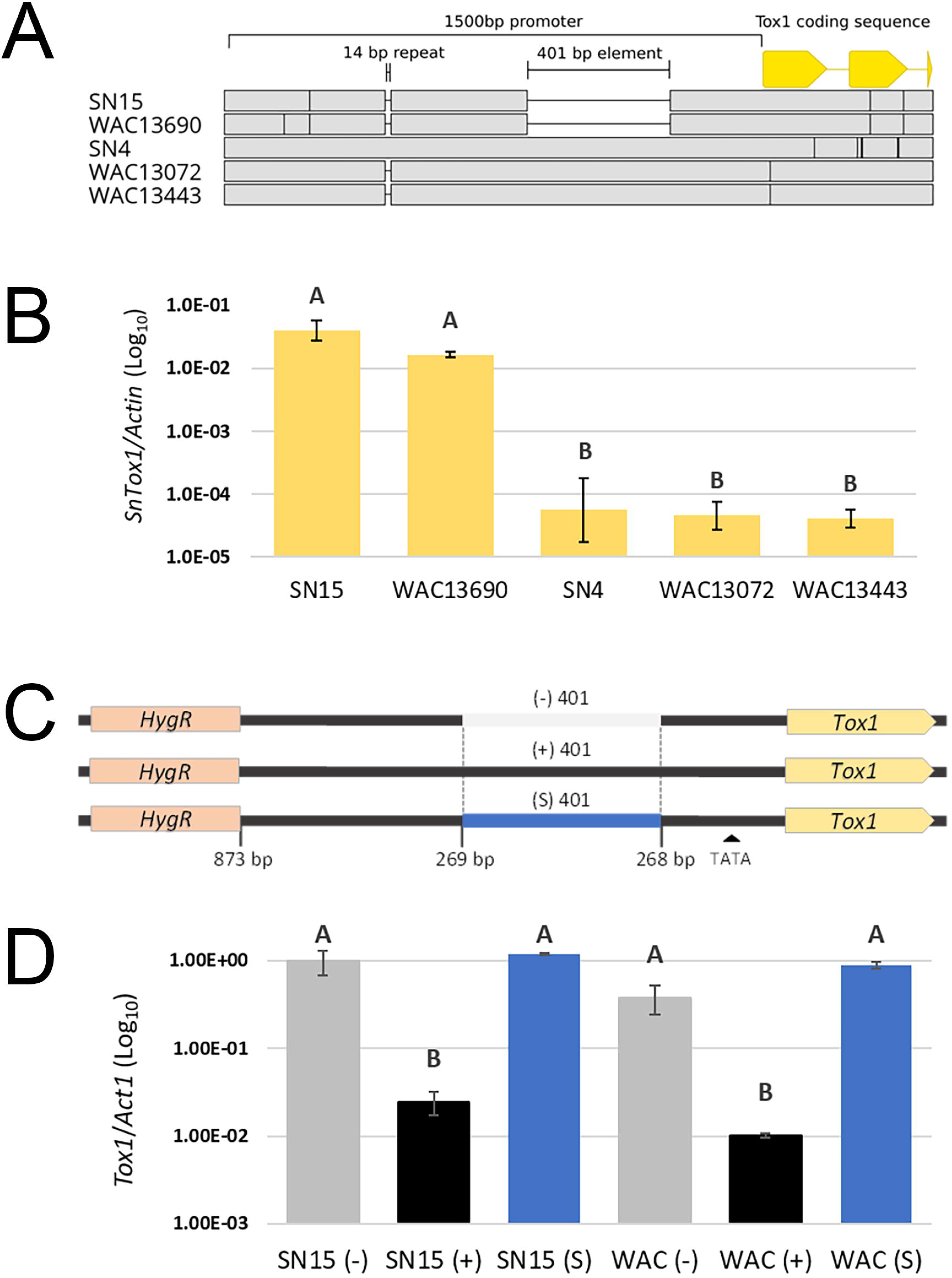
Sequence specificity of the 401 bp PE401 dictates *Tox1* repression. A. Alignment of the promoter regions comparing the USA reference isolate SN4 with four Australian isolates +/- the 401 bp PE401. B. *Tox1* expression [2^dCt(*Tox1-Actin*)^] of *P. nodorum* wildtype isolates +/- PE401 *in vitro*. Error bars indicate standard deviations from biological triplicates. Letters not connected by the same letter are significantly different (*P* < 0.05) based on ANOVA. C. Schematic overview of the three *Tox1* promoter replacement strategies with (+), without (-) or using a spacer (S) 401 bp sequence into SN15 and WAC13443 (WAC) strains (S2 File). D. *Tox1* expression (2^dCt(*Tox1-Actin*)^) in SN15 and WAC mutant strains carrying promoter replacements grown *in vitro*. Error bars indicate standard deviations from combined averages (*n* = 2) of independently verified isogenic promoter replacement mutants. Letters not connected by the same letter are significantly different (*P* < 0.05) based on ANOVA.

We then created isogenic strains with and without PE401 to eliminate the possibility of strain-specific genetic factors located outside of the *Tox1* locus involved in gene repression. The constructs used for promoter replacement included 873 bp of the *Tox1* promoter either with PE401 (+), without PE401 (-) or with a 401 bp spacer (S) (Fig 3C). These were independently transformed into SN15 and WAC13443 at the *Tox1* locus through homologous recombination. Quantitative RT-PCR analysis of the mutants revealed a 40-fold (SN15) and 38-fold (WAC13443) reduction in *Tox1* gene expression occurred in the presence of PE401 when grown in Fries3 broth *in vitro*. *Tox1* was not repressed in mutants that carry the spacer sequence, suggesting the effect was attributable to sequence specificity conferred by PE401 (Fig 3D). We concluded that PE401 is associated with *Tox1* repression, perhaps through an interaction with a trans-acting regulatory factor.

We then determined the regulatory role of PE401 in *Tox1* expression during host infection using qRT-PCR. Infection time points were chosen to represent major stages of infection such as penetration (3 dpi), colonisation (5 dpi) and sporulation (7 dpi) (46) (Fig 4). As expected, *Tox1* expression was much lower (log power) in SN15 and WAC13443 mutants carrying PE401 than mutants carrying either the spacer sequence or lacking PE401 throughout the entire infection period (Fig 4).

**Fig 4.**
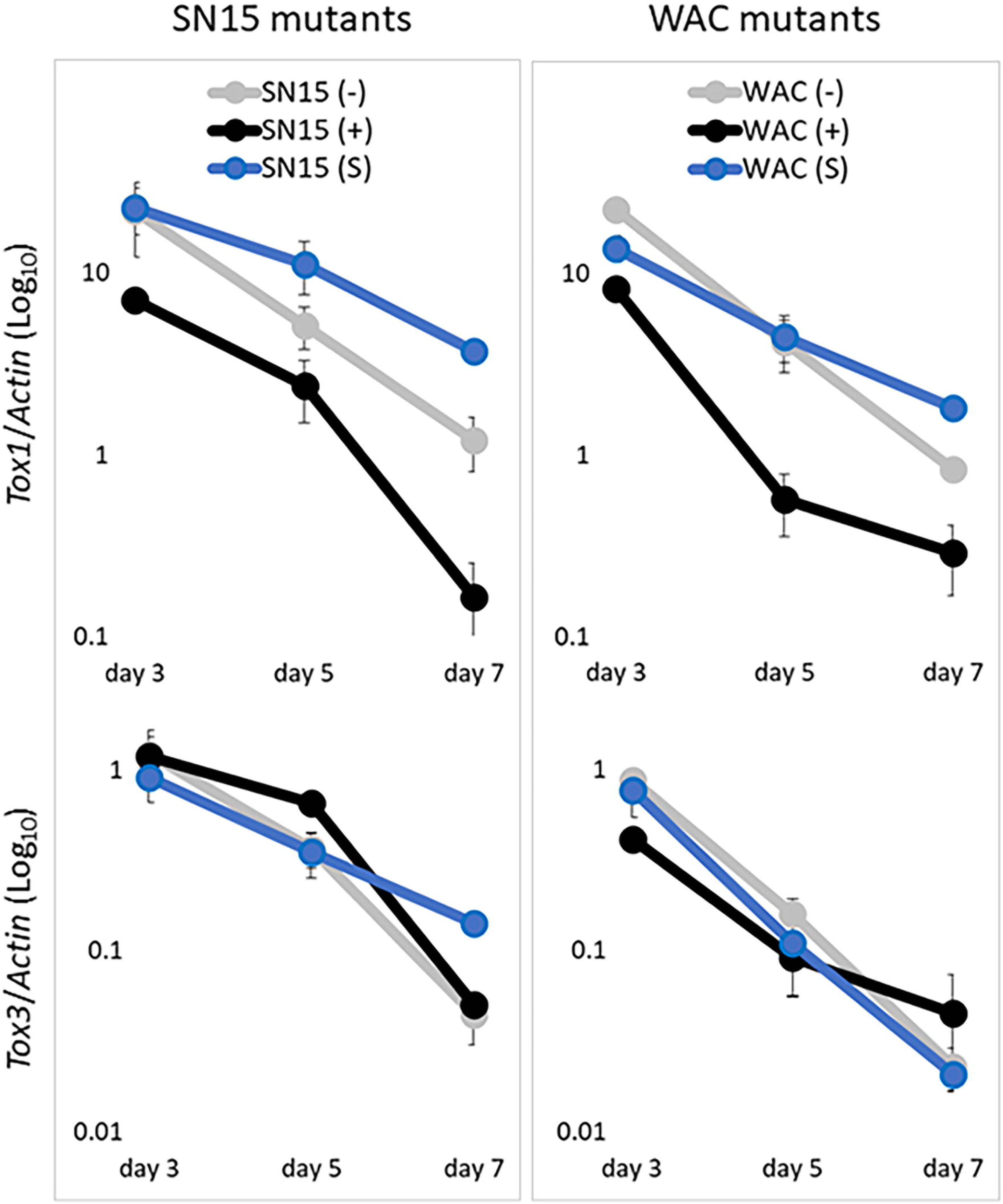
PE401 represses *Tox1* expression during host infection. Quantitative RT-PCR determination of *Tox1* and *Tox3* expression in promoter replacement mutants of SN15 and WAC13343 sampled at three (early penetration), five (colonisation) and seven (sporulation) days post-infection of wheat cv. Halberd. The average values [2^dCt(*Target-Actin*)^] of two biological replicates were used. Error bars indicate standard deviations.

It was previously observed that deletion of *Tox1* in SN15 caused a significant increase in *Tox3* expression (22). Therefore, we hypothesised that the inclusion of PE401 would alleviate *Tox3* repression by *Tox1*. However, the decrease in *Tox1* expression conferred by PE401 did not affect *Tox3* expression (Fig 4). This indicated *Tox1* repression caused by PE401 was insufficient to alleviate epistasis of *Tox3* and that full suppression of *Tox1* expression is needed to achieve this.

### PE401 influences Tox1-*Snn1*-mediated ETS in wheat

The contribution of Tox1-*Snn1* to SNB symptoms caused by SN15 (+) and SN15 (-) mutant strains were determined using an association mapping approach. We used a Calingiri (*Snn1*, *snn3b1*, *tsn1*) and Wyalkatchem (*snn1*, S*nn3B1*, *tsn1*) (CxW) DH wheat population infected with mutant spores at the seedling stage to directly compare the effect of PE401 repression on Tox1-*Snn1* and other important SNB interactions. SNB seedling resistance QTL can be detected in the CxW population on chromosomes 1BS (*Snn1*), 5BS (*Snn3B1*), 2A, 2D, 3A and 4B when infected with SN15 or its derivative NE deletion mutants (22). The seedling infection assay revealed the average disease score on the 179 wheat lines (including parental lines) for the SN15 (+) mutant was 4.4. This is significantly lower than SN15 (-) at 4.8 (Table 2). We then compared the SNB severity of SN15 (+) and (-) on genotype combinations that possess *Snn1* and observed that the former isolates were significantly more pathogenic (Table 2). In contrast, no significant difference was observed between SN15 (-) and SN15 (+) on *snn1* wheat lines.

**Table 2.**
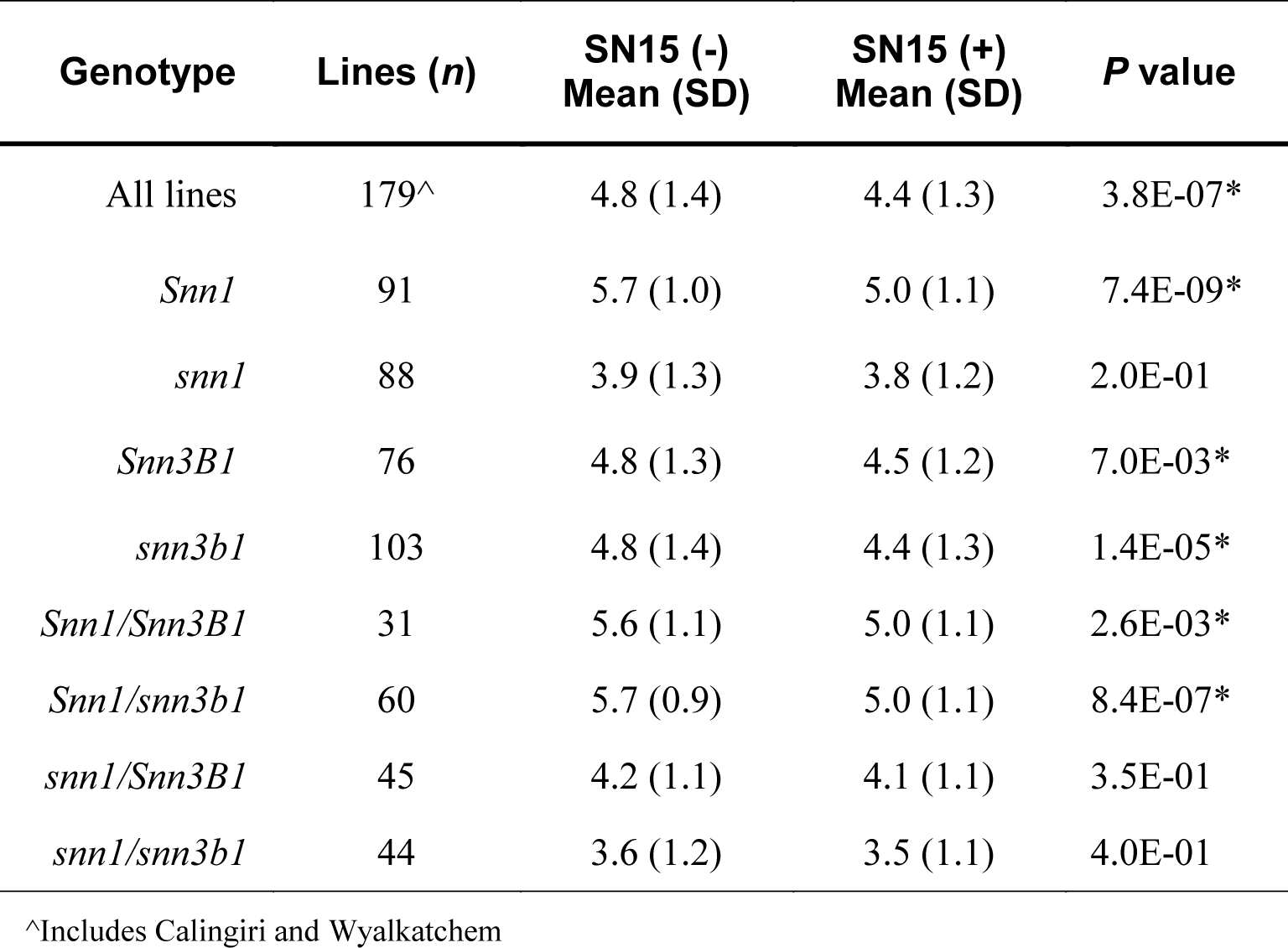
Seedling SNB rating of the CxW DH population grouped by Tox1 and Tox3 sensitivity conferred by Snn1 and Snn3B1, respectively. SN15 isogenic mutants with (+) and without (-) PE401 were used to assess seedling wheat infection. Mean SNB scores and SD were determined for both promoter replacement mutants. A paired Student’s *T*-test (*n* = 3) was used to compare the mean between strains for each wheat genotype groups. *Indicates significant differences between the mutants at the *P*<0.05 significance threshold. Disease scores for all lines are provided in S3 File.

There was no difference in disease severity on *Snn3* versus *snn3* wheat for either mutant, which had previously been observed for the *tox1* mutant compared to SN15 (22). This indicated that *Tox1* remained epistatic to *Tox3* when expressed at the lower level conferred by the PE401. Reliable markers for the 2A, 2D, 3A and 4B resistance loci are not yet available which meant the disease severity could not be directly compared between the SN15 (-) and (+) mutants. Nevertheless, the results suggest that the Tox1-*Snn1* interaction contributed significantly to SNB severity, and that higher expression of *Tox1* conferred by the absence of PE401 produced a significantly higher level of disease on *Snn1* wheat.

### The absence of PE401 results in higher biomass on *Snn1* wheat lines

We hypothesised that *P. nodorum* isolates lacking PE401 resulting in higher *Tox1* expression are fitter on *Snn1* wheats. However, low *Tox1*-expressing *P. nodorum* isolates that possess PE401 are maintained in the population pool at a low frequency (S1 Fig, 47). To determine evidence of fitness penalty on *Snn1* wheat, we used a digital PCR approach to compare the DNA biomass of SN15 (+) and (-) during infection of *Snn1* (H086, H336 and H091) and *snn1* (H213 and H324) wheat varieties deriving from the CxW population that lacked Tox3 and Tox2A sensitivity (Fig 5). Fungal biomass was determined at two and four dpi where *Tox1* is maximally expressed. For *snn1* wheat varieties infected with either SN15 (+) or (-), there was no significant difference (*P* > 0.05) between the biomass of SN15 (+) and (-) at 2 or 4 dpi, with an expected increase of biomass over time. For wheat varieties with *Snn1*, the biomass of SN15 (-) was significantly higher than SN15 (+) at two and four dpi (Fig 5). We then co-infected all five wheat varieties with an equal ratio of SN15 (+) and (-) to determine if SN15 (-) would outcompete SN15 (+) during infection on *Snn1* wheats using a basic De Wit replacement series of 50:50 input replacement combining SN15 (-) and (+) (48). We observed the proportion of SN15 (+) and (-) biomass was near-identical at both time points in contrast to the initial hypothesis. This indicates that SN15 (-) assisted SN15 (+) infection by driving Tox1-*Snn1* ETS of the host through elevated *Tox1* expression. Moreover, in two *Snn1* wheat lines (H086 and H336) the total biomass at four dpi of co-infection was slightly but significantly elevated (*P* < 0.05) compared to SN15 (-) (Fig 5). This suggests that SN15 (-) assists SN15 (+) in host colonisation possibly through a mutualistic interaction.

**Fig 5.**
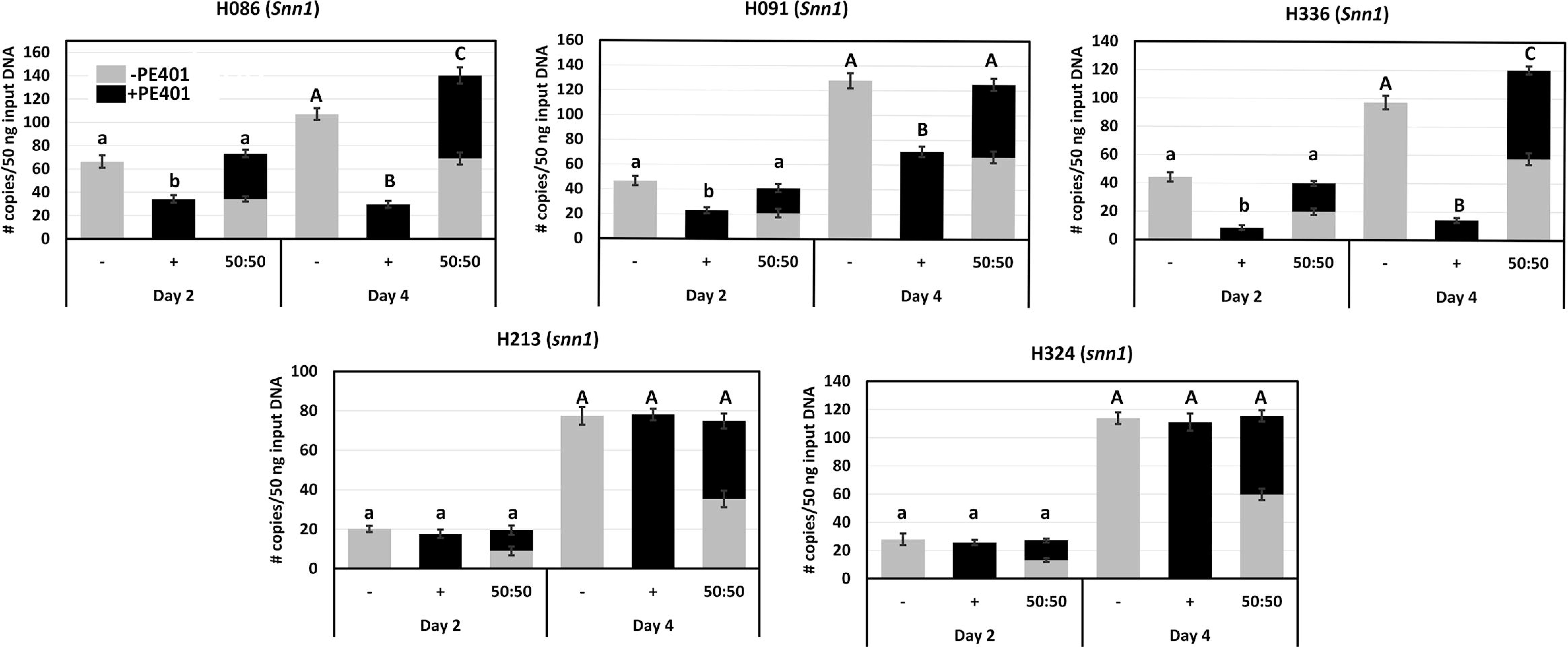
Digital droplet PCR analysis of SN15 (+) and (-) biomass during infection of wheat varieties with and without *Snn1* (bar graph) at 2 and 4 dpi where *Tox1* expression is maximal. Relative biomass was also determined on a 50:50 co-infection of SN15 (+) and (-) (stacking bar graph). All lines selected were insensitive to Tox3 (*snn3b1*) and *P. nodorum toxa13* culture filtrate containing Tox2A (22). A one-way ANOVA with posthoc Tukey-HSD testing was used to identify significant (*P* < 0.05) differences across treatments of two (lowercase letter) and four (uppercase letter). Letters not connected by the same letter are significantly different.

### *Tox1* repression by PE401 alleviates epistasis of a major SNB QTL on chromosome 2A

*Tox1* is epistatic to the SNB QTL *Qsnb.cur–2AS1* detected on chromosome 2A in addition to Tox3-*Snn3B1* (22). Since neither the NE or host receptor on chromosome 2A have been cloned, epistasis can only be studied through genetic mapping using wheat mapping populations that segregate for *Qsnb.cur–2AS1*. Therefore, interval mapping and QTL analysis were undertaken using the markers previously developed for the CxW population (22) to explore any differences in the disease interactions produced by the SN15 (-) and (+) PE401 mutant strains. The QTL detected on chromosome 1B, where *Snn1* is located, explained the highest phenotypic contribution to SNB in the SN15 (-) mutant (47.9%) compared with SN15 (+) (30.1%) (Table 3). However, the *Qsnb.cur–2AS1* SNB QTL was detected in the SN15 (+) mutant only, where it contributed 12.1% to the disease. *Qsnb.cur–2AS1* was the most prominent QTL previously detected during infection with the *Tox1* knockout mutant *tox1-6* (S2 Fig) (22). Despite Wyalkatchem being a donor of *Snn3B1* in the CxW population (22), association mapping did not detect the *Snn3B1* QTL on chromosome 5B. Therefore, we concluded that *Tox1* repression by PE401 alleviated the epistatic effect on Tox2A-*Qsnb.cur– 2AS1* but not Tox3-*Snn3B1*.

**Table 3.**
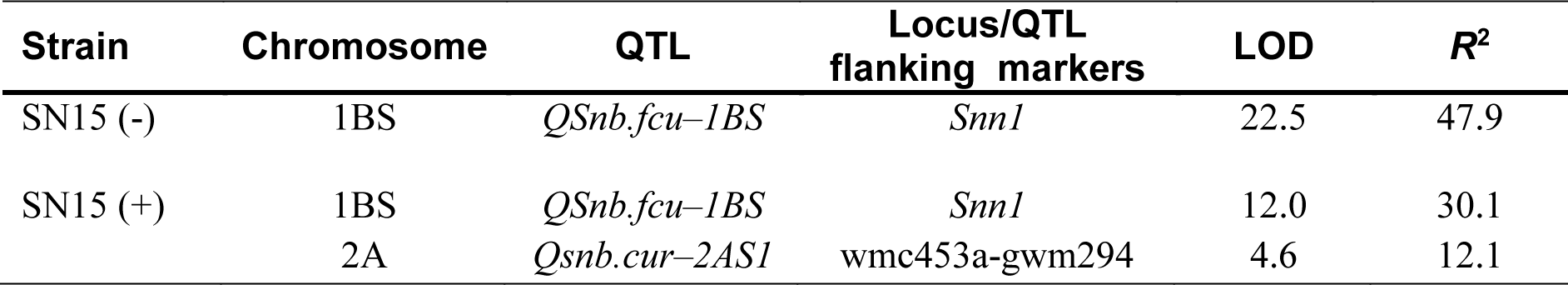
A summary of SNB QTL detected with the CxW wheat population. Details of the flanking markers, LOD scores and phenotype contribution (*R*^2^) are indicated. Composite interval mapping of SNB QTL is provided in S2 Fig.

### Promoter substitution identifies the positive regulatory region of *Tox1*

To characterise the core regulatory regions involved in gene expression, we generated SN15 promoter replacement mutants that carry sequentially truncated *Tox1* promoter sequences (Fig 6). The shortest truncation included only the *Tox1* 5’ UTR (untranslated region) from the annotated genome which encompassed 86 bp upstream of the *Tox1* start codon (35). Further truncations were made at 107 bp and 157 bp upstream of the ATG, either side of a TATATAA sequence typical of a core eukaryotic TATA box element (49). Additional truncations were made at 263 bp and 310 bp upstream to exclude/include the site of the PE401 variant. In addition, 598 bp and 873 bp upstream mutants were selected to exclude possible undiscovered regulatory elements; the 873 bp region corresponding to the entire sequence of the SN15 (-) mutant used in the preceding analyses. Furthermore, SN15 (+) mutants carrying PE401 were generated in each of the 310, 598 and 873 bp mutant backgrounds to assess any effect of the upstream elements on the repressor activity (Fig 6A).

**Fig 6.**
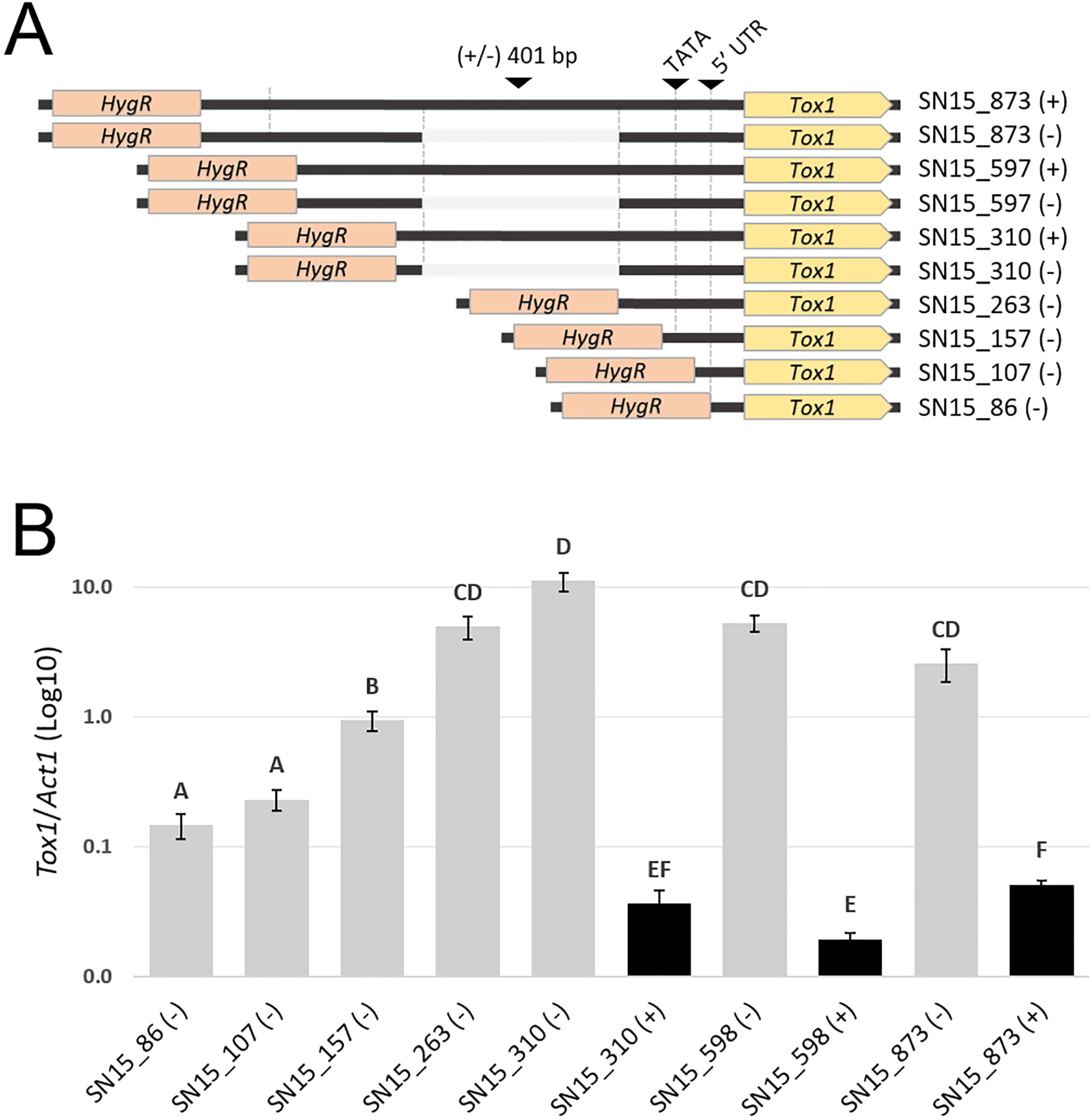
Analysis of *Tox1* expression regulation through promoter sequential deletions. A. Schematic overview of the *Tox1* sequential promoter deletions in SN15 either with (+) or without (-) the 401 bp PE401. Indicated are the predicted TATA box and 5’ UTR derived from the reference SN15 genome. B. *Tox1* gene expression [2^dCt(*Tox1-Actin*)^] in the corresponding SN15 mutants after 72 hrs growth in Fries3 liquid culture. Error bars indicate standard deviations from the combined averages two biological replicate of two independently verified isogenic promoter replacement mutants. Letters not connected by the same letter are significantly different (P < 0.05) based on ANOVA.

Gene expression analysis revealed that rather than any single element being essential, an additive effect on *Tox1* gene expression was observed by the inclusion of regions up to 310 bp, where the expression was maximal (Fig 6B). In particular, significant increases were observed by the inclusion of the putative TATA box region (107 - 157 bp) and the upstream region (157 – 263 bp). PE401 acted as a strong repressor in each of the promoter deletion mutants tested. The inclusion of longer promoter regions did not significantly increase *Tox1* gene expression. This suggested that positive regulatory elements are located directly adjacent to the 3’ end of the PE401 which functions as a repressor sequence independent of upstream sequences.

## Discussion

In this study, we sought to characterise the gene regulatory elements controlling *Tox1* expression and explore their role in NE epistasis and the virulence of *P*. *nodorum*. An important finding was the clear role in *Tox1* repression conferred by PE401 located 267 bp upstream of the start codon of *Tox1* (Fig 6). Analysis of the truncated promoter mutants indicated PE401 could repress the transcriptional activators driving *Tox1* expression (Fig 6). The exact mechanism remains obscure, as there were no distinctive features of a non-coding element or MGE. Nonetheless, we can conclude that a novel polymorphic NE regulatory element has been identified in *P*. *nodorum*, likely targeted by sequence-specific repressor proteins.

The most parsimonious explanation regarding the origins of PE401 is that the sequence was present in the ancestral *Tox1* gene promoter and was subsequently lost in a subset of isolates. This stems from the fact that it was only detected in isolates outside the proposed centres of origin for SNB (44, 45). Populations carrying the highest proportion of isolates without PE401 were from South Africa and Australia. There is little data regarding South African wheat but it is known that *Snn1* wheat in Australia has been widely sown throughout the period corresponding to the isolate collections assessed (24, 47). No detectable shifts in the frequency of the element were observed during this time 2001 and 2015 (S1 Fig). This indicates either that selection was neutral throughout this period, or that the maintenance of some variation in the population is beneficial for the pathogen. This situation is supported by the results of our co-infection assay, which demonstrated PE401 did not cause a fitness penalty in the presence of the strain lacking PE401. The other major *P*. *nodorum* population where PE401 was absent is the North Dakota population, albeit at a reduced frequency (27%) compared to Australia (87%) and South Africa (100%). In this region, it has been reported that *Snn1* is prevalent in widely grown durum wheat varieties (16, 50). Interestingly, *P*. *nodorum* population structure analyses have suggested the North Dakota population is unrelated to those in Australia and South Africa, which are closely related to each other (44, 51). It is therefore possible that PE401 was lost on separate occasions in these populations. On the other hand, it is also possible that genetic exchange has occurred between the populations followed by selection. Accordingly, if *Snn1* is removed from cultivars sown in these and other regions, it will be important to monitor the frequency of the element in addition to the *Tox1* gene in fungal isolates.

The infection assay using the CxW DH population demonstrated that higher expression of *Tox1* conferred a higher average level of virulence on *Snn1* wheat. While it is known that a fungal NE repertoire dictates pathogenic fitness, the role of gene expression had not been extensively examined (52). In the case of the tan spot of wheat fungus *Pyrenophora tritici-repentis*, a much higher level of *ToxA* expression than in *P. nodorum* is maintained during infection (53). It was proposed this can compensate for the reduced potency of the PtrToxA isoform relative to *P*. *nodorum* variants (53) and demonstrates an example in another pathosystem where NE expression change can have significant disease outcomes.

The genetic mapping results comparing SN15 (-) and SN15 (+) on the CxW DH wheat population suggested removal of PE401 did not affect Tox1-*Snn1* epistasis over Tox3-*Snn3B1*. This indicated that Tox1-*Snn1* must entirely be removed before Tox3-*Snn3B1* compensates for virulence. However, the *Qsnb.cur–2AS1* SNB QTL was only detected by the inclusion of PE401. The physical interval for *Qsnb.cur–2AS1* is notably large (22). Nevertheless, several recent studies suggest the locus contains a gene conferring sensitivity to an undiscovered NE, that functions as an important determinant of both seedling, adult leaf and glume SNB resistance since these QTL overlap with *Qsnb.cur–2AS1* (20, 22, 26, 54). In addition, a minor seedling resistance QTL on chromosome 2A was detected in a recombinant inbred population derived from Swiss winter wheat varieties Arina and Forno (20). Therefore, the Tox2A- *Qsnb.cur–2AS1* interaction provides a layer of NE redundancy that maintains virulence on wheat, albeit at a lower level, in the absence of other interactions such as Tox1-*Snn1*. In addition to SNB, a co-localised *Qsnb.cur-2AS1* QTL is a major contributor to tan spot resistance in wheat (55). Interestingly, tan spot is the most damaging necrotrophic fungal disease of wheat in Australia (56) and the fungus frequently co-infects wheat with *P. nodorum* (57–59).

Regional *P. nodorum* populations differ greatly in NE gene frequency (60, 61), possess high haplotype variability (45), differential isoform activity (52), variable NE expression (62; this study) and high evolutionary potential (1). This makes SNB a difficult disease to eliminate using a common genetic resistance background in wheat across different cereal growing regions where SNB is endemic.

PE401 provides a novel example of a population variant regulating NE epistasis through gene repression. Epistasis between NE-host gene interactions was frequently observed through the use of association mapping with wheat genetic mapping populations. A complex picture has now emerged as some interactions are additive in SNB (23) while others are epistatic (Table 1). Several instances of epistasis, where one fungal NE-host receptor interaction suppresses the disease contribution of another, are now documented (Table 1). In some cases, differential NE gene expression underpins epistasis, but the regulatory elements were unclear. In one example, the expression of *ToxA* was reported to be two-fold higher during infection in the *P. nodorum* isolate SN5 compared with SN4 (62). Higher ToxA expression correlated with a greater contribution of ToxA-*Tsn1* to the disease phenotype of SN5 on the BR34 x Grandin RI wheat population that segregates for *Tsn1* and *Snn2*. Conversely, the phenotypic contribution of Tox2-*Snn2* was reduced (62). Recently, a gene encoding the NE Tox267, which interacts with *Snn2*, has been cloned and functionally characterised (17). The deletion of *Tox267* in SN4 resulted in up-regulation of the NEs *ToxA*, *Tox1*, and *Tox3*, demonstrating further connectivity between the expression of these genes. Another study that used an RI wheat population (ITMI) segregating for *Tsn1*, *Snn1* and *Snn3* reported that the presence of *ToxA* in SN2000 suppressed Tox1-*Snn1* ETS (31). Gene deletion of *ToxA* produced only a mild reduction in disease severity, which was compensated by increased *Tox1* expression. Collectively, these studies suggest gene repression as the basis for NE epistasis is a common mechanism, for which the results presented here have provided a novel mechanistic insight.

The phenomenon of NE epistasis is also observed in other fungal-plant pathosystems, dictating the contribution of effector-susceptibility/resistance gene interactions during disease (27). *P. tritici-repentis* possesses a near-identical copy of *ToxA* to *P. nodorum* and infects the host through *Tsn1*-mediated ETS (13). The deletion of *ToxA* increased the virulence of *P. tritici-repentis* on some *Tsn1* wheat lines and unmasked the effect of a novel chlorosis-inducing factor (63, 64). In the Brassicae black leg *Leptosphaeria maculans*-*Brassica napus* pathosystem, *Rlm3*-mediated resistance through recognition of the avirulence effector protein AvrLm3, was suppressed by another avirulence effector AvrLm4-7 (65).

Further evidence was provided here using a competitive De Wit replacement assay that demonstrated a direct selective advantage in fungal proliferation in the absence of the PE401 on *Snn1* wheat lines since the high-expressing *Tox1* variant confers a virulence advantage over low *Tox1* expression. In Australia, *Snn1* is continually being maintained in wheat varieties grown in Western Australia (WA), a region where SNB is endemic (47). Currently, wheat cv Scepter that lacks *Snn1* is the most widely grown in WA (52% area sown and increasing) (66, www.caigeproject.org.au/germplasm-evaluation/bread/disease-screening/toxicity/). Varieties with *Snn1* are currently sown at very low percentages (66). Maintenance of high *Tox1* expression can be bioenergetically taxing if it does not confer a selective advantage for the pathogen. Therefore, it remains to be seen if the decline in *Snn1* frequency will affect the frequency of PE401 within the Australian *P. nodorum* population over the next few years through selection pressure incurred by the mass adoption of Scepter. Tox1 and the other two well-characterised NEs ToxA and Tox3 are present in almost all *P. nodorum* isolates in Australia but in other parts of the world, *ToxA*, *1* and *3* frequencies are much lower (45, 60). Coupled with the presence of other NE-*Snn* interactions that may be active and epistatic to Tox1-*Snn1* (Table 1), we can begin to speculate that there is less dependency for Tox1-*Snn1* ETS outside of Australia. This could explain why PE401 is being retained at a high frequency in *P. nodorum* isolates in the Northern Hemisphere. It was also hypothesised by Haugrud *et al*.. (30) that *P. nodorum* down-regulates *Tox1* as a means to prevent antagonism with the activity of other NEs that subvert alternative host-immune responses. This hypothesis requires further testing.

From the co-infection assay, we anticipated that SN15 (-) will outcompete SN15 (+) when infecting *Snn1* wheats when all other known NE-host dominant susceptibility gene interactions are removed in CxW DH lines. In contrast, SN15 (-) appears to ‘rescue’ SN15(+) when co-infecting. In two of the three *Snn1* wheat lines, the pathogen complex accumulated more biomass than a single infection. McDonald *et al.* (67) observed different genotypes of *P. nodorum* on a single infected wheat plant which suggests that *P. nodorum* exists as a fungal disease complex to ensure maximal infection capabilities on susceptible wheat. This observation accounts for a mixed *Tox1* expressing genotype in the Australian *P. nodorum* population where the high expressing variant remains dominant (47, S1 Fig).

In this study, we observed that *Tox1* exhibits both ‘selfish’ and ‘selfless’ (altruistic) characteristics. We hypothesise that the loss of PE401 allow the gene to exert dominance or epistasis over both Tox3-*Snn3* and Tox2A-*Qsnb.cur-2AS1* interactions which is driven by *Snn1* and results in *Tox1* exhibiting a ‘selfish’ character (68). However, our coinfection competition assay demonstrates evidence of altruism at the isolate level where one isolate must have a higher *Tox1* expressing variant to help maintain the virulence and prevalence of isolates with lower *Tox1* expression, which is mediated by the presence of PE401 in the *P. nodorum* population. Hence, this study has provided a unique insight into a ’NE arms race’ that is occurring within the *P*. *nodorum* population.

The identity of trans-acting regulatory molecules that are associated with *Tox1* regulation remains unknown at this stage. However, key results from this study allow us to propose a regulatory model for *Tox1* and NE-*Snn* epistasis detected using the CxW population (Fig 7). In the absence of the PE401, transcriptional activator(s) can drive high *Tox1* expression (Fig 7A). As a result, Tox1-*Snn1* contributes strongly to SNB while the *Qsnb.cur-2AS1* QTL is suppressed. An unidentified repressor binds to the PE401 and inhibits the transcriptional activator(s) bound to the TATA-region (69, 70), thereby repressing *Tox1* expression and the consequent epistatic effect on the Tox2A-*Qsnb.cur-2AS1* interaction during SNB (Fig 7B). Although the proposed regulatory mechanism remains speculative at this stage, it provides a framework that will allow future studies of regulatory components. As such, affinity purification assays using these sequences as bait represent a promising avenue to identify the regulators involved.

**Fig 7.**
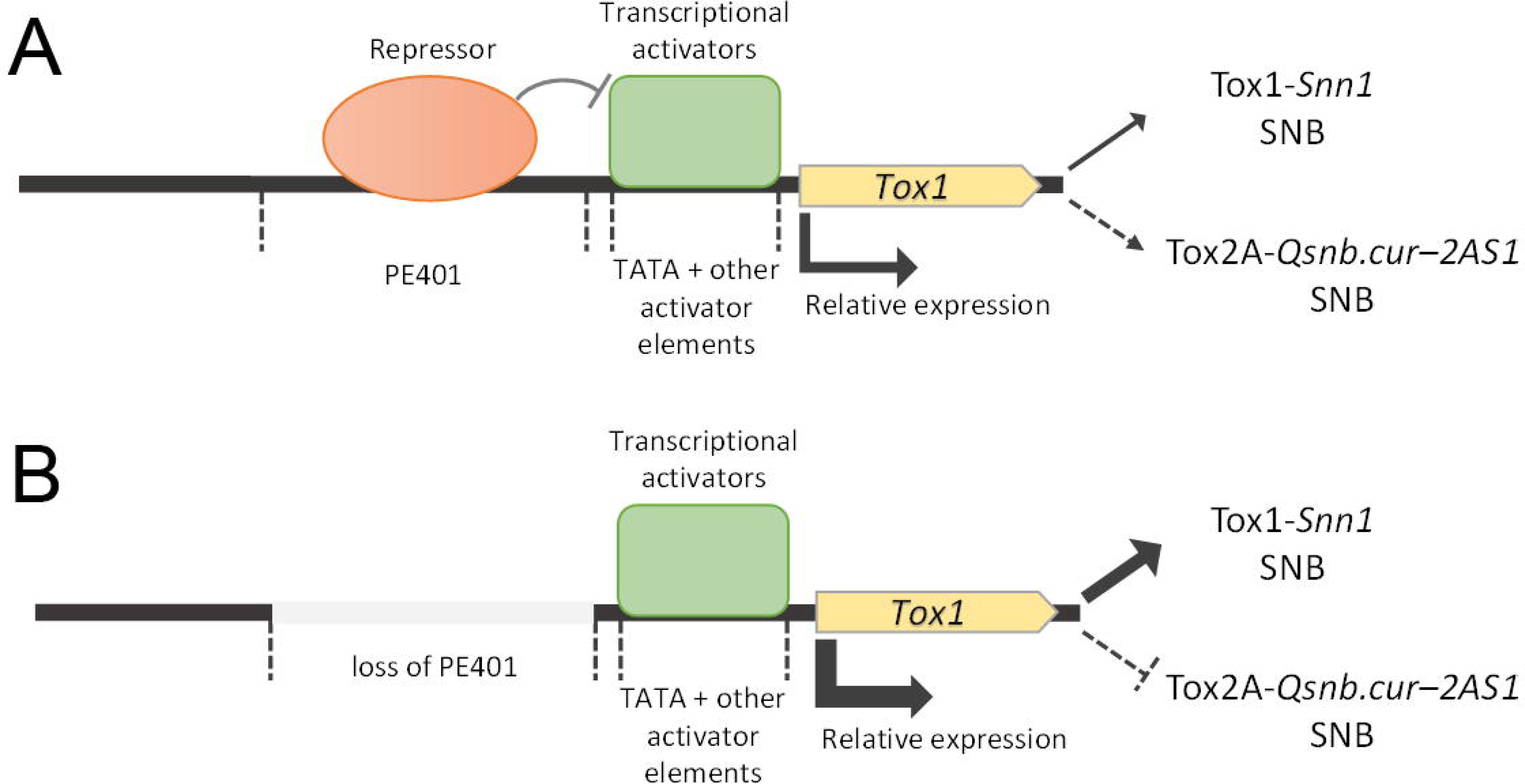
The proposed model for *Tox1* regulation and its influence on SNB. SNB contribution A. in the absence and B. presence of the 401 bp PE401. The effect on gene expression and direct (solid arrows) or epistatic (dashed arrows) contribution to SNB are depicted by the strength of the arrows.

### Conclusions and implications

The findings presented here have advanced our knowledge regarding the control of *Tox1* expression in *P. nodorum* and its profound epistatic effect on SNB resistance in wheat. Although the proposed regulatory mechanism remains speculative at this stage, we have defined individual regulatory elements within the promoter region of *Tox1*. This provides a framework for future studies into the components regulating *Tox1*. We are currently using a DNA-protein bioassay approach to identify transcriptional activator and repressor proteins that are associated with the promoter region of *Tox1* and regulate its expression. However, underlying questions on the mechanism of the epistatic suppression of NE gene expression and phenotypic contributions of SNB remained unanswered. Small interfering RNA may be present within the fungus that regulates gene expression including silencing NE genes (71). Alternatively, NE-host gene interactions within the same system compete for common plant-signaling pathways, such as the mitogen-activated protein kinase pathway, or are antagonistic in the mechanisms by which ETS is triggered (15, 72).

The evidence presented highlights the general importance of understanding NE regulation, the outcomes of which extend beyond the individual NE-host receptor interactions. The evolution of changes in NE gene expression provides a mechanism for *P*. *nodorum* to adapt where the complement of host receptors shifts through resistance breeding efforts. More importantly, the identification of PE401 will help to formulate effective crop protection strategies against SNB of wheat. Downie *et al.* (23) provided a framework of management strategies to minimise the impact of SNB. This includes establishing an extensive isolate collection programme, epidemic monitoring, population genotyping and wheat variety adoption strategies. Based on the outcome of this study, we propose additional guidelines for SNB management to complement the Downie *et al.* (23) framework. These are;

1. Monitor for changes in the frequency of PE401 in regional *P. nodorum* populations.
2. The maintenance of *Snn1* in wheat is a likely driver for *P. nodorum* to retain PE401 and *Tox1* evolution. Therefore, *Snn1* removal should be prioritised in cereal breeding programmes through conventional breeding, genome editing or genetic modifications.
3. Introduce the resistance allele for *Qsnb.cur-2AS1* with other SNB resistance alleles (eg. *snn1* conferring Tox1 insensitivity) into wheat breeding stocks. Phan *et al.* (9) and Lin *et al.* (54) demonstrated that stacking of four regionally relevant SNB resistance alleles provide optimal disease resistance in wheat.
4. *Qsnb.cur-2AS1* also confers resistance to tan spot (55). Inclusion of the *Qsnb.cur-2AS1* insensitivity/SNB resistance allele in breeding programmes should reduce the impact of the SNB-tan spot disease complex.

We anticipate that the outcome of this study will drive a greater level of research into the field of effector regulation and epistasis in other fungal-plant pathosystems to generate similar outcomes to improve existing crop protection strategies.

## Materials and methods

### Fungal culture

All *P. nodorum* strains were maintained on V8-PDA agar at 21°C under a 12-h photoperiod or grown in Fries3 broth as previously described (53, 73).

### Compilation of *Tox1* promoter variants

The *P. nodorum Tox1* sequence was derived from the genome annotation SNOG_20078 of the Australian reference isolate SN15 (35). A BLAST database was built (Geneious Prime 20.2.5) from the other available published and assembled genomes for 33 *P. nodorum* and *Parastagonospora avenae* f. sp. *tritici* isolates (36, 37). The corresponding *Tox1* loci, including the 1,500 bp upstream regions, were retrieved using the SN15 *Tox1* nucleotide sequence. The 24 sequences retrieved were aligned using the Geneious aligner on the default settings to detect polymorphisms in the promoter region. 159 Australian isolates were then PCR screened (Tox1_screen_F/R) to detect product size shifts corresponding to a 401 bp element presence/absence variant (47). All primer sequences are shown in S1 Table. Sanger sequencing was undertaken on the region for two Australian isolates with the 401 bp element (WAC13443 and WAC13072) and an additional isolate without (WAC13690). The *Tox1* locus and 1,500 bp upstream regions were analysed for a worldwide set of 146 *P. nodorum* isolates that were derived from Illumina genome assemblies published previously (51, 74). The NCBI BLAST service (75) was then used to query the sequence read archives for 184 Illumina sequenced genomes for USA isolates deposited as PRJNA398070 (76), which were non-redundant to the isolates already compiled. The *Tox1* nucleotide sequence and 1000 bp promoter from SN4 was used to query the respective archives for each isolate. Where continuous coverage of reads >95% identity was obtained across the corresponding region, the presence/absence of PE401 was able to be determined. Isolate details used for promoter alignment are described in S1 File. ***In silico* analyses of the *Tox1* promoter variants**

A world distribution map was produced from the geographical information assembled for a panel of 489 *P. nodorum* isolates with or without the *Tox1* PE401 (S1 File) using ggmaps (77). The SN4 *Tox1* promoter sequence was then queried for matches in Repbase by submitting the 1500 bp region through the CENSOR portal using the default setting (78, 79). The 401 bp SN4 *Tox1* PE401 was also submitted to the Rfam portal (80). Furthermore, the NCBI BLAST service (75) was used to scan the PE401 sequence against the SN15 genome, the NCBI nucleotide and the whole-genome shotgun collections. Dot plots and GC% graphs were produced using Geneious. The SNOG_30065 expression and gene annotation data were obtained from a previous study (42).

### Generating *Tox1* promoter replacement mutants

The cloning strategy and an overview of the modified *Tox1* promoter locus in fungal transformants are depicted in Fig 8 and the primer details are listed in S1 Table. In summary, promoter replacement constructs were assembled in two stages by attaching flanking regions to a *pTef1-HygR-tTef1* resistance marker using a Golden Gate’ (GG) style cloning system (81) established in-house using a pUC19 vector backbone (New England Biolabs, Ipswich, Massachusetts, USA) and kindly provided by Dr. Jordi Muria-Gonzales. In stage one the *pTef1- HygR-tTef1* marker was assembled by BbsI-mediated cloning using *Tef1* promoter and terminator regions derived from *Aspergillus nidulans* using pTef1_P_BbsI_F/R and tTef1_T_BbsI_F/R, as well as the hygromycin resistance gene amplified from Pan7 (Reference = https://link.springer.com/article/10.1007/BF00279797) with HygR_B_BbsI_F/R. Stage two involved the assembly and attachment of distinct flanks to the marker cassette using *Bsa*I cloning. The left flank was the same for all constructs, amplified from SN15 using pTox1_HR_FL_Bsa1_F/R. The right flanks were amplified from SN15 gDNA for the (-) PE401 construct variant using pTox1_873_HR_FR_Bsa1_F/pTox1_HR_FR_Bsa1_R. The same primers were used in combination with pTox1_Dom_BsaI_F/R to amplify the right flank from WAC13443 gDNA for PE401 constructs (which allowed fragment domestication for GG cloning by the introduction of a point mutation). PE401 was also substituted in the previous construct by amplifying the GG plasmid with pTox1_indel_3_BbsI_F/pTox1_indel_5_BbsI_R and ligating in a 401 bp spacer (S) element with a similar GC content but an unrelated sequence (S2 File). This was amplified from the pGEM®-T-Easy vector (Promega, Madison, WI) using the primer pair spacer_BbsI_F/R. The three resulting homologous recombination (HR) constructs, were amplified and purified from these templates using pTox1_HR_FL_F/R, before polyethylene glycol (PEG) mediated transformation into SN15 and WAC13443. Truncated versions of the *Tox1* promoter replacement construct were produced by modifying the forward primer used to amplify the right flank to incorporate either 86, 107, 157, 263, 310 (+/- PE401) or 598 bp (+/- PE401) regions upstream of the translational start site. The resulting constructs were PEG-transformed into SN15 as previously described (82). All cloned constructs were verified through PCR screening and Sanger sequencing, while the fungal transformants were PCR screened using pTox1_HR_screen_F/tTox1_HR_screen_R and by qPCR (Tox1_qPCR_F/R vs Actin_qPCR_F/R) to confirm single-copy integration using a robust quantitative PCR method (83). Two single copy mutants for all transformants were retained for gene expression analyses. All genetically modified strains and their background strains are described in Table 4.

**Fig 8.**
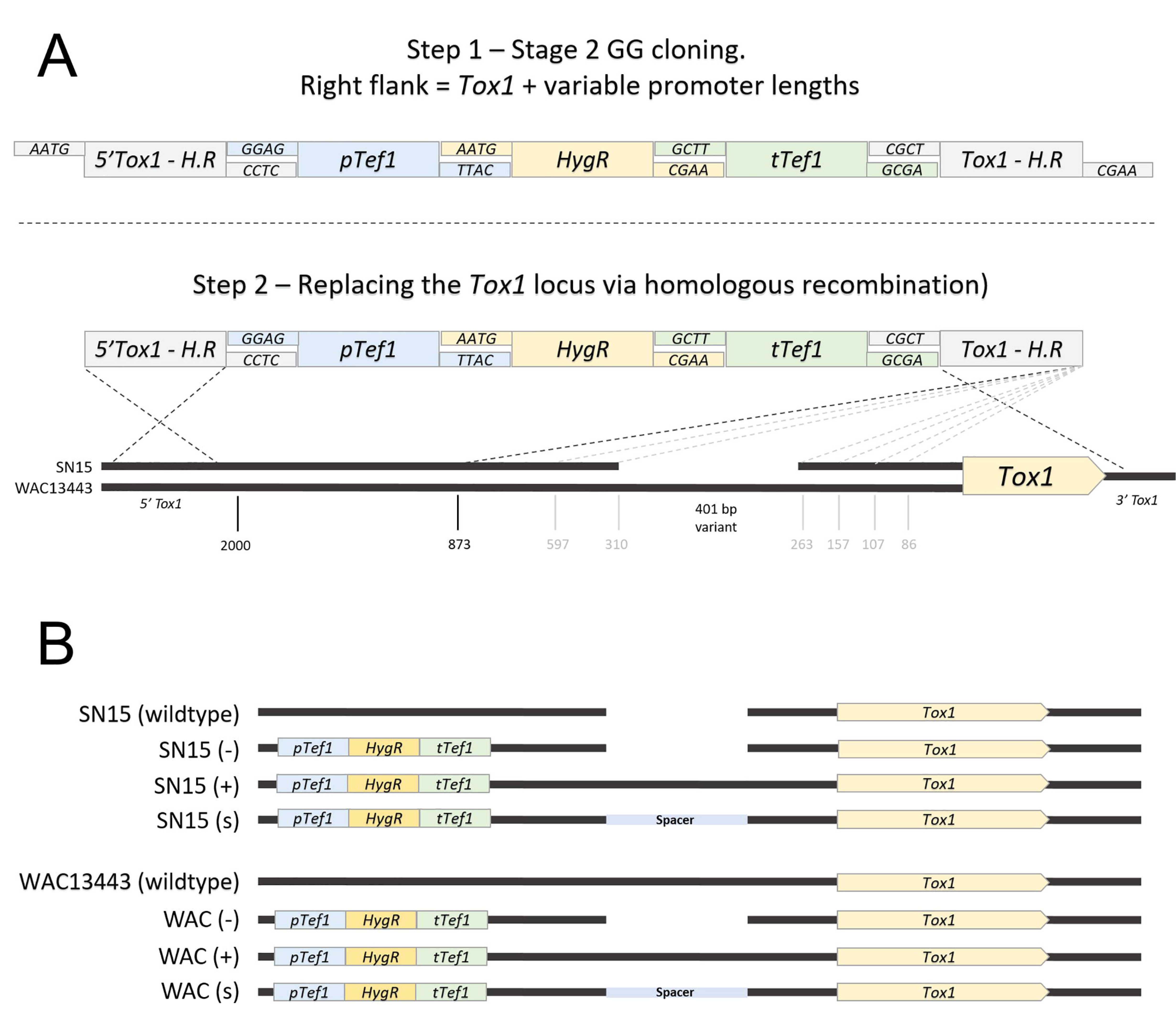
*Tox1* promoter replacement strategy. A. In step 1, Golden Gate cloning was used to assemble constructs for promoter replacement at the *Tox1* locus. Linear replacement constructs were amplified for fungal transformation in step 2. B. Promoter replacement mutants generated in the study representing the *Tox1* locus in the respective background strains.

**Table 4.** Fungal mutant strains used in this study. Wildtype strains used for mutagenesis are indicated.

| Strain | Description | Source |
| --- | --- | --- |
| SN15 | *Parastagonospora nodorum* wild-type. | *DPIRD |
| WAC13443 | *Parastagonospora nodorum* wild-type. | *DPIRD |
| SN15 (-) | SN15 transformed without PE401. | This study |
| SN15 (+) | SN15 transformed with PE401. | This study |
| SN15 (S) | SN15 transformed with a spacer sequence in place of PE401. | This study |
| WAC (-) | WAC13443 transformed without PE401. | This study |
| WAC (+) | WAC13443 transformed with PE401. | This study |
| WAC (S) | WAC13443 transformed with a spacer sequence in place of PE401. | This study |
| SN15_86 (-) | SN15 expressing *Tox1* with a promoter truncated at 86 bp upstream of ATG. -PE401. | This study |
| SN15_107 (-) | SN15 expressing *Tox1* with a promoter truncated at 107 bp upstream of ATG. - PE401. | This study |
| SN15_157 (-) | SN15 expressing *Tox1* with a promoter truncated at 157 bp upstream of ATG. - PE401. | This study |
| SN15_263 (-) | SN15 expressing *Tox1* with a promoter truncated at 263 bp upstream of ATG. -PE401. | This study |
| SN15_310 (-) | SN15 expressing *Tox1* with a promoter truncated at 310 bp upstream of ATG. -PE401. | This study |
| SN15_310 (+) | SN15 expressing *Tox1* with a promoter truncated at 310 bp upstream of ATG. +PE401. | This study |
| SN15_597 (-) | SN15 expressing *Tox1* with a promoter truncated at 597 bp upstream of ATG. - PE401. | This study |
| SN15_597 (+) | SN15 expressing *Tox1* with a promoter truncated at 597 bp upstream of ATG. +PE401. | This study |
| SN15_873 (-) | SN15 expressing *Tox1* with a promoter truncated at 873 bp upstream of ATG. -PE401. | This study |
| SN15_873 (+) | SN15 expressing *Tox1* with a promoter truncated at 873 bp upstream of ATG. +PE401. | This study |

\*Department of Primary Industries and Regional Development

### Gene expression analysis

*P. nodorum* cDNA was synthesised from the respective strains/mutants grown for three days in sterile Fries3 liquid medium for three days shaken at 120 rpm in the dark as previously described (53). Quantitative RT-PCR was used to determine the expression level of *Tox1* normalised to *Act1* (2^dCt^). For *in planta* time series analysis of gene expression, lesions were harvested at 3, 5 and 7 days post-infection by the seedling spray method, from which cDNA was synthesised for qPCR as previously described (84). Tox1_qPCR_F/R and Tox3_qPCR_F/R were used to assess the respective gene expression normalised to *Act1* (2^dCt^). A one-way ANOVA with Tukey-HSD post-hoc test was used to test for differences (p<0.05) between isolates and/or mutants (SPSS version 27.0).

### Association mapping and QTL analysis

The CxW DH wheat population that consisted of 177 lines (Intergrain Pty Ltd, Perth, Australia) was used to carry out association mapping and QTL studies of seedling SNB as previously described (22). A total of 482 DarT, wheat genomic SSR and EST-SSR markers were used to construct a genetic map for the population (22). Two-week old seedlings were infected with *P. nodorum* using a whole plant spray assay was visually assessed for SNB severity on a standardised scale of one to nine as previously described (22). A score of one indicates no disease symptoms whereas a score of nine indicates a fully necrotised plant (9, 22). QTL mapping was undertaken using MultiQTL v. 2.6-Complete software (MultiQTL Ltd, Institute of Evolution, Haifa University, Israel). The QTL mapping was based on a genetic linkage map previously built for the CxW population using the Kosambi mapping function using MultiPoint v. 3.2 (MultiQTL Ltd, Haifa University, Israel) from maximum recombination frequencies of 0.35 (22). This included 385 markers polymorphic between the CxW parent lines. Seedling disease scores taken from the average of three randomised biological replicates were used for interval mapping to determine QTL linked to SNB. Markers with the logarithm of the odds (LOD) score set at a ≥ 2.5 cut-off was used to construct an interval model for the corresponding QTL as previously described (22).

### Attached leaf infection assay

An attached leaf assay with modifications was used to carry out multi-strain co-infection (85). Briefly, the first leaves of two-week old wheat seedlings were inoculated with a *P. nodorum* spore suspension [1 x 10^6^ spores/ml in 0.02 % (v/v) Tween 20] using an airbrush kit (6ml/min, Ozito, Perth, Australia). Per treatment (+PE401; -PE401; 50:50 mix) a total of 6 ml spore inoculum was evenly sprayed across five wheat lines with 6 seedlings per line. Tween was used as a negative control. The wheat lines were H213, H324 (*snn1*, *snn3b1*, insensitive to Tox2A) and H086, H336, H091 (*Snn1*, *snn3b1*, insensitive to Tox2A). Leaves were allowed to dry and incubated in a controlled growth chamber as described previously (85). Two and four days post-inoculation, infected leaves were harvested with a scalpel and snap-frozen in liquid nitrogen (three biological repeats per time point per wheat line per treatment). DNA was extracted using phenol/chloroform method as described previously (86) and concentrations were measured using a Qubit Fluorometer (ThermoFisher Scientific, MA USA). All samples were diluted to a concentration of 10 ng/ul DNA.

### Droplet digital PCR quantification for fungal biomass quantification

Droplet digital PCR (ddPCR) was used for absolute quantification of biomass base on fungal DNA copy abundance. This approach allows for the precise and absolute quantification of DNA biomass as it measures and analyses about 20,000 reactions (droplets) in a single well (87). We used two primer sets, a specific SN15 (+) primer targeting PE401 (pTox1_401_qPCR_F and pTox1_401_qPCR_R) and a general SN15 primer (ddPCRf and ddPCRr) that will detect both SN15 (-) and SN15 (+) DNA (S1 Table). In a single well, all the positive droplets carrying a copy of the target sequence are counted and expressed as the number of copies per input DNA concentration (in our case, 50 ng of total DNA). As such, we determined and compared the number of copies between the treatments and wheat lines as an accurate measure of pathogen biomass. We used the Evagreen (BioRad, IL USA) approach as described previously (88). Briefly, a 96-well plate was loaded into the QX200 Auto DG to generate droplets (Bio-Rad, IL USA) in each well. Next, plates were sealed using the PX1 PCR Plate Sealer before continuing to PCR (C1000 Touch Thermal Cycler, Bio-Rad, IL USA) which was performed with the primer sets described above using the following program: 95°C for 5 min, 44 cycles of 95°C for 30 s, 63°C for 30 s, 72°C for 30 s and finish with 4°C for 5 min followed by 90°C for 5 min. The plate was subsequently loaded into the QX200 Plate Reader (Bio-Rad, IL USA) and data was analysed using the QuantaSoft Software (Bio-Rad, IL USA). Genomic DNA of SN15 (-) and SN15 (+) strains were included as positive controls and additionally, primer efficiency was tested, thereby ensuring comparability between wells. A one-way ANOVA with posthoc Tukey-HSD testing was used to identify significant (*P* < 0.05) differences between the three infection treatments (+401 bp; -401 bp; 50:50 mix) per wheat line (SPSS version 27.0).

## Acknowledgements

This study was supported by the Centre for Crop and Disease Management, a joint initiative of Curtin University and the Grains Research and Development Corporation (research grant CUR00023 Project F3). We thank Intergrain Pty. Ltd. for the wheat mapping population. We thank Mr. Lincoln Harper and Dr. Noel Knight for the technical advice on ddPCR. EJ was supported by the Australian Government Research Training Program Scholarship.

## Author contributions

KCT conceived the experiment. KCT, EJ and SJ carried out experimental designs. EJ, SJ and HP conducted the experiments. KCT, KS and RO provided project supervision. LL conducted critical preliminary experiments. DC performed analyses and provided datasets. DP provided datasets. KCT and EJ wrote the manuscript. All authors edited the manuscript.

## Supporting information caption

**S1 File.** *Tox1* isolate metadata describing the isolate ID, species, country, region and host information and collection date. The presence of the 401 bp PE401 in the *Tox1* promoter is indicated along with the data source and reference from which this was determined.

**S2 File.** PE401 and the 401 bp spacer sequence used to substitute the element in SN15 (S) control mutants. A dotplot is provided to demonstrate the sequence dissimilarity.

**S3 File.** SNB disease score and QTL mapping data. A table indicating the CxW lines tested, the presence of *Snn1* and *Snn3*-linked marker, the average disease scores for SN15 (-) and SN15 (+) mutants as well as a summary of the disease score statistical analysis and the chromosomal linkage groups analysed.

**S1 Fig.** Temporal assessment of 401 bp PE401 frequencies in the Australian *P. nodorum* isolate collection.

**S2 Fig.** Composite interval mapping of QTL associated with SNB caused by SN15 (-), SN15 (+), SN15 and *tox1-6* (22).

**S1 Table.** Primers used in this study.

## Notes

### Competing Interest Statement

The authors have declared no competing interest.

