## Supplementary material for "Variability in an effector gene promoter of a necrotrophic fungal pathogen dictates epistasis and effector-triggered susceptibility in wheat": S2 File

>401 bp element

ACGCTGTACGCTATACGCAGACTGTTAGGAGATCACACCTTATATATGCTGTTTTACTCCTTTACTCTATAGTCCTATCAATCGTGTACTTAGTCGAGAGTTCTAGTATATTAGGTTCGCAACTTACATTAGGAAGGCGACTAATCATACTGTAAGGTGAAATTGGCAGAAGCCTTCAGGAAAATCTCTAGATACCTATGGTAGCTCTGAAGGAAGCTTACAGCCGTTTGCTGACAGGTTCGAGCACATACTACAGACATCTAATCTTAGCCAATATTCTGCCCCTGCGTCAGCATTAAGGACTTCCAATTAATTAGTCTGTTCTAGGAAACCTCAGAGCTTCCCGCAAATCCAAGGAATATAGTACAACAGCAGAGAGGCCTCTGAGACCTATTCTTAAT

>401 bp spacer

GCTTGCTCCTTTCGCTTTCTTCCCTTCCTTTCTCGCCACGTTCGCCGGCTTTCCCCGTCAAGCTCTAAATCGGGGGCTCCCTTTAGGGTTCCGATTTAGTGCTTTACGGCACCTCGACCCCAAAAAACTTGATTAGGGTGATGGTTCACGTAGTGGGCCATCGCCCTGATAGACGGTTTTTCGCCCTTTGACGTTGGAGTCCACGTTCTTTAATAGTGGACTCTTGTTCCAAACTGGAACAACACTCAACCCTATCTCGGTCTATTCTTTTGATTTATAAGGGATTTTGCCGATTTCGGCCTATTGGTTAAAAAATGAGCTGATTTAACAAAAATTTAACGCGAATTTTAACAAAATATTAACGTTTACAATTTCAGGTGGCACTTTTCGGGGAAATAATG

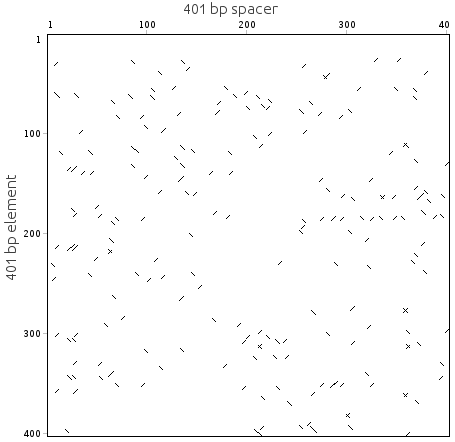
