## Supplementary figures and images for "Variability in an effector gene promoter of a necrotrophic fungal pathogen dictates epistasis and effector-triggered susceptibility in wheat"

### S1 Fig

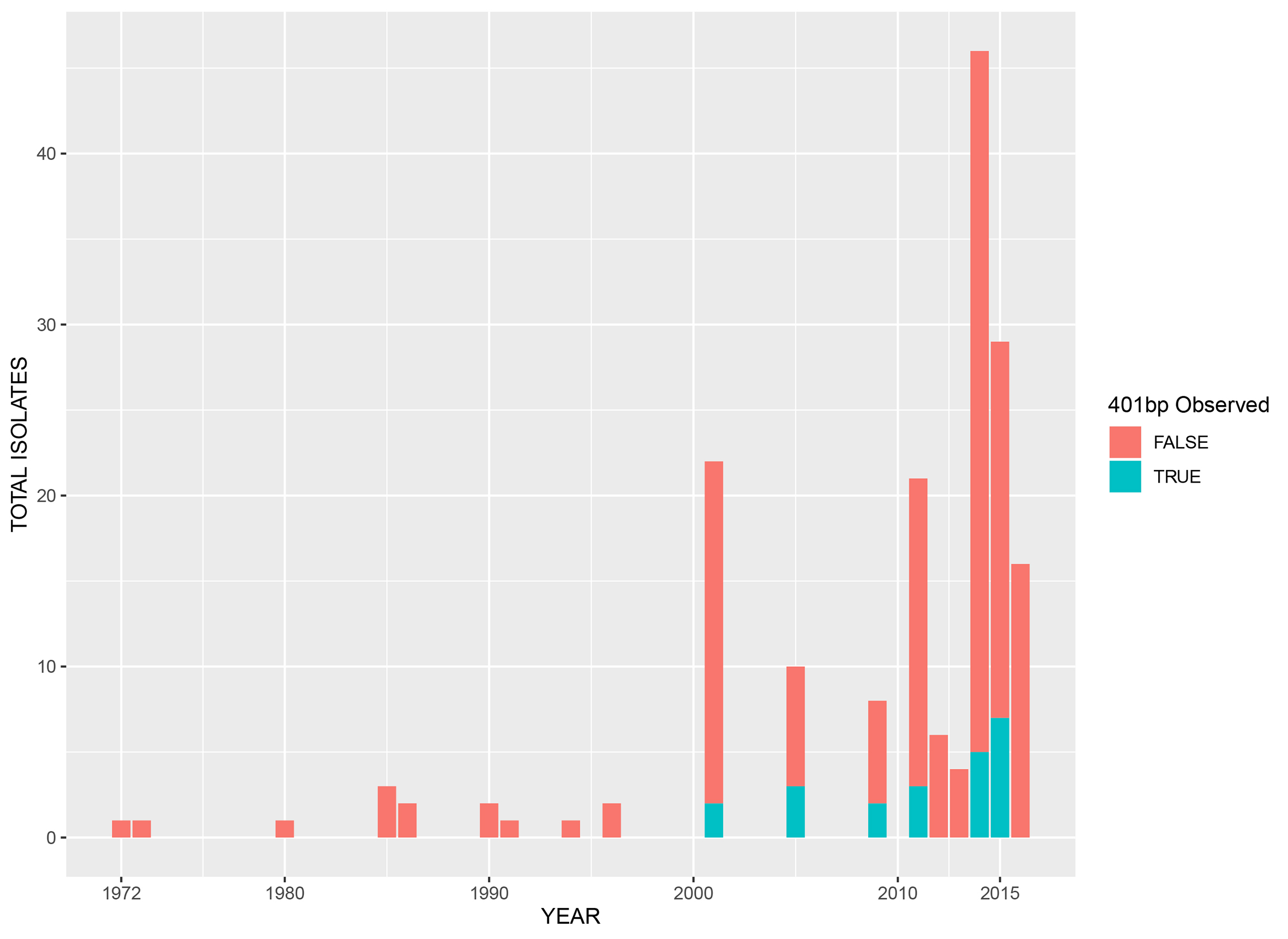

### S2 Fig

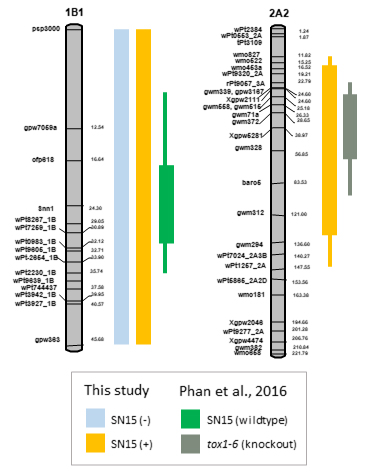
